# Measurement of Vaccine Direct Effects under the Test-Negative Design

**DOI:** 10.1101/237503

**Authors:** Joseph A. Lewnard, Christine Tedijanto, Benjamin J. Cowling, Marc Lipsitch

**Affiliations:** Division of Epidemiology, School of Public Health, University of California, Berkeley, Berkeley, California; Center for Communicable Disease Dynamics, Department of Epidemiology, Harvard TH Chan School of Public Health, Boston, Massachusetts; WHO Collaborating Center for Infectious Disease Epidemiology and Control, School of Public Health, Li Ka Shing Faculty of Medicine, The University of Hong Kong, Hong Kong Special Administrative Region, China.

**Keywords:** Test-negative design, influenza, vaccine effectiveness

## Abstract

Test-negative designs are commonplace in assessments of influenza vaccination effectiveness, estimating this value from the exposure odds ratio (OR) of vaccination among individuals treated for acute respiratory illness who test positive for influenza virus infection. This approach is widely believed to recover the vaccine direct effect by correcting for differential healthcare-seeking behavior among vaccinated and unvaccinated persons. However, the relation of the measured OR to true vaccine effectiveness is poorly understood. We derive the OR under circumstances of real-world test-negative studies. The OR recovers the vaccine direct effect when two conditions are met: (1) individuals’ vaccination decisions are uncorrelated with exposure or susceptibility to the test-positive or test-negative conditions, and (2) vaccination confers “all-or-nothing” protection (whereby certain individuals have no protection while others are perfectly protected). Biased effect size estimates arise if either condition is unmet. Such bias may suggest misleading associations of vaccine effectiveness with time since vaccination or the force of infection of influenza. The test-negative design may also fail to correct for differential healthcare-seeking behavior among vaccinated and unvaccinated persons without stringent criteria for enrollment and testing. Our findings demonstrate a need to reassess how data from test-negative studies can inform policy decisions.

---

Observational study designs (1,2) are needed to measure vaccine effectiveness (VE) when randomized trials are infeasible or unethical, as with the new formulations of influenza vaccines used each year (3). The “test-negative” design—a modification of the traditional case-control design—has become popular for measuring clinical effectiveness of seasonal influenza vaccines (1). It resembles earlier designs such as the indirect cohort method (4) and the selection of “imitation disease” controls in case-control studies (5). Individuals who experience acute respiratory illness (ARI) and present for care receive a laboratory test for influenza virus infection, and their vaccination history is ascertained. The exposure-odds ratio of vaccination among test-positive and test-negative subjects (OR), in some instances adjusted for potential confounding using stratification or regression, has frequently been used to measure VE (6), where 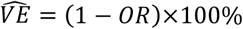 (7). Causal interpretations of resulting estimates have become the basis for policymaking, such as the US Advisory Committee on Immunization Practices recommendation that quadrivalent live attenuated influenza vaccine (LAIV) should not be used in the US during the 2016-17 and 2017-18 seasons (3,8,9).

Unlike VE estimates from traditional case-control studies, the test-negative measure is expected to correct for differential treatment-seeking behaviors among vaccinated and unvaccinated persons because only individuals who seek care are included (10). However, potential confounding, misclassification, and selection biases under the test-negative design (2,11–13) have ignited debate about the suitability of test-negative studies as a basis for policymaking. Whereas directed acyclic graphs have been useful in revealing such biases (9,14,15), quantitative implications of these biases for VE estimates remain uncertain (16).

To resolve this uncertainty, we derived the relation of the test-negative OR to true VE, defined as the vaccine-conferred reduction in susceptibility to influenza infection and/or influenza-caused ARI (vaccine “direct effect” (17)). We used this mathematical relationship to assess the quantitative impact of potential biases in test-negative studies. We consider a test-negative study of VE against seasonal influenza as a guiding example, noting that our findings have implications for test-negative studies of vaccines against rotavirus (18,19), cholera (20,21), meningococcus (22), pneumococcus (4), and other infections.

## NOTATION

For consistency, we use notation from a previous study (10) where possible; we list all parameters and definitions in **Table 1**. Assume that ARI may result from influenza infection (*I*) or other causes (*N*). Susceptible individuals acquire infection at time-constant rates *λ_I_* and *λ_N_*; we show later that results hold for seasonal or otherwise time-varying acquisition rates *λ*(*t*). We define *t*=0 as the start of the influenza season, and assume individuals are vaccinated around this time (before extensive transmission). Infections cause ARI with probability *π_I_* and *π_N_*, respectively. Out of the entire population *P*, a proportion of individuals (*v*) received vaccine. Because individuals who opted for vaccination may differ from others in their likelihood for seeking treatment for ARI, define the probability of seeking treatment for an ARI episode as *μ_V_* among the vaccinated and *μ_U_* among the unvaccinated; we address how differential treatment seeking for test-positive and test-negative conditions influences estimates in a later section.

**Table 1:**
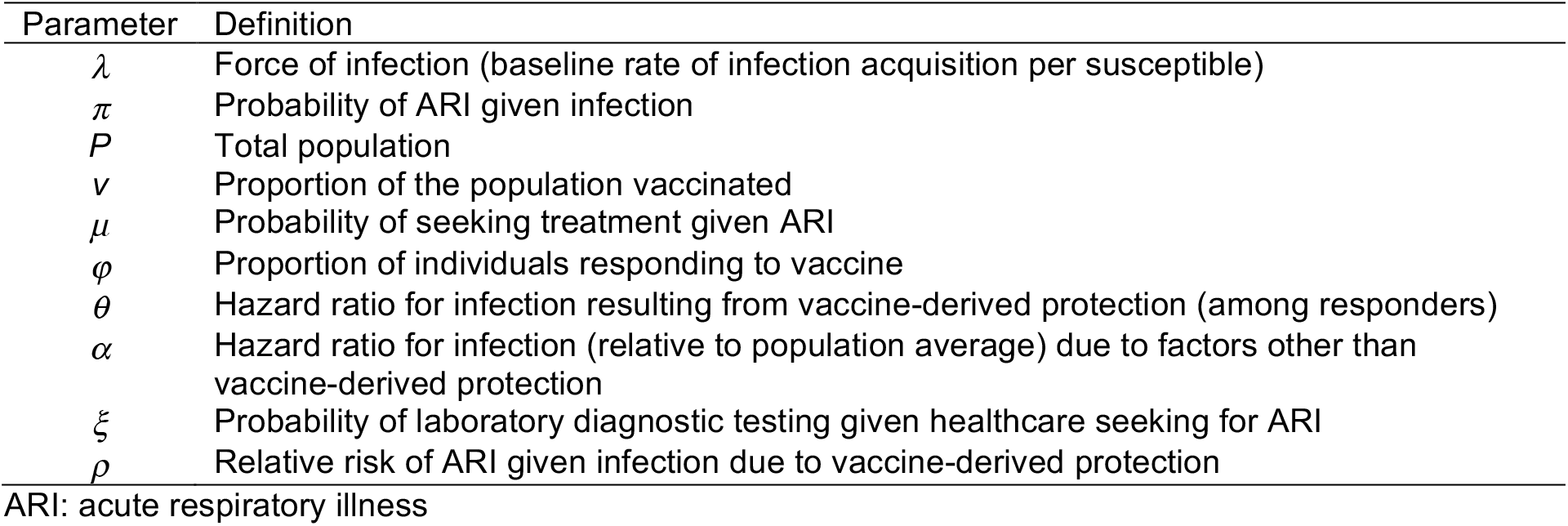
Parameters (referenced in order of appearance)

| Parameter | Definition |
| --- | --- |
| $\lambda$ | Force of infection (baseline rate of infection acquisition per susceptible) |
| $\pi$ | Probability of ARI given infection |
| $P$ | Total population |
| $\nu$ | Proportion of the population vaccinated |
| $\mu$ | Probability of seeking treatment given ARI |
| $\varphi$ | Proportion of individuals responding to vaccine |
| $\theta$ | Hazard ratio for infection resulting from vaccine-derived protection (among responders) |
| $\alpha$ | Hazard ratio for infection (relative to population average) due to factors other than vaccine-derived protection |
| $\xi$ | Probability of laboratory diagnostic testing given healthcare seeking for ARI |
| $\rho$ | Relative risk of ARI given infection due to vaccine-derived protection |

ARI: acute respiratory illness

Because a single type or subtype of influenza typically dominates each season, assume that naturally-acquired immunity protects against within-season re-acquisition of influenza. The proportion of individuals remaining susceptible to infection at time *t* is thus *e^−λ_I_t^*. Assume further that the various non-influenza causes of ARI (*N*) are unlikely to provide immunity against one another, so that the full population remains at risk of *N* throughout; we show later that this assumption does not impact estimates (**Web Appendix 1**).

Consider two mechanisms by which vaccination protects against infection. Define *φ* as the proportion of individuals responding to vaccine, so that a proportion 1 − *φ* remain unaffected by vaccination; here we assume individuals’ likelihood of responding is unassociated with exposure or susceptibility to infection. Among the responders, define *θ* as the hazard ratio for infection (measured relative to the hazard rate of infection among non-responders and unvaccinated persons) resulting from vaccine-derived protection (23,24). The special case where *θ*=0 and 0<*φ*<1 corresponds to a situation of “all-or-nothing” protection for responders and non-responders, respectively, while “leaky” protection for all recipients arises under *φ*=1 and 0<*θ*<1 (23,24,17), whereby all vaccine recipients experience a reduced rate of acquiring infection. We note that this definition of “leaky” protection is unrelated to the relative risk for vaccine recipients and non-recipients to experience progression of infection to symptomatic disease (17), and consider this issue in a subsequent section. The general circumstances of 0<φ<1 and 0<*θ*<1 correspond to an intermediate scenario of “leaky-or-nothing” protection. Perfect protection attains for *θ*=0 and *φ*=1, and no protection attains when *φ*=0 (no individuals respond to vaccination) or *θ*=1 (responders receive no protection). The vaccine direct effect on susceptibility to infection is the rate ratio of infection given vaccination:

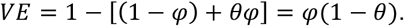

This parameter is of interest in vaccine studies as the basis for calculating the effective reproductive number and the critical population to vaccinate (25). To highlight design-level features most pertinent to the interpretation of test-negative studies, and in line with typical reporting of VE estimates, our analysis does not address heterogeneity in vaccine response beyond the consideration of “all-or-nothing” and “leaky-or-nothing” protection, nor do we address impacts of vaccination on infectiousness, as estimates from conventional test-negative studies do not capture indirect effects. We refer readers to previous studies addressing such issues in the contexts of differing study designs (17,26–28). Where applicable, we address sources of confounding in test-negative studies that may lead to incorrect inferences of heterogeneity in vaccine effects among individuals or over time.

## PERFORMANCE OF THE ODDS RATIO UNDER VACCINATION UNCONFOUNDED BY EXPOSURE OR SUSCEPTIBILITY TO THE CONDITIONS

Here we consider the case where individuals’ decision-making about whether to receive influenza vaccine is uncorrelated with their *a priori* risk of acquiring influenza and test-negative conditions, and with the probability that these conditions would cause ARI (or another clinical endpoint of interest for study enrollment; *π_I_* and *π_N_*). To examine the potential for the test-negative design to correct for treatment-seeking biases, we allow vaccine recipients and non-recipients to have different probabilities of seeking treatment for ARI (*μ_V_* and *μ_U_*), assuming for now that these probabilities are unaffected by the cause of the ARI. We relax this assumption in a later section.

To understand what the OR measures in test-negative studies, we derive the rate at which individuals enter into the study as test-positive or test-negative subjects given their vaccination status. The rate of ascertaining test-positive, vaccinated persons is

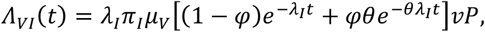

where the force of infection (*λ_I_*) is applied upon as-yet uninfected members of the vaccinated population; we further account for the proportion (*π_I_μ_V_*) of individuals expected to show symptoms and seek treatment. The rate of ascertaining test-positive, unvaccinated subjects is

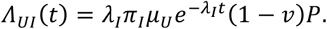

Test-negative vaccinated and unvaccinated persons are ascertained at the rates

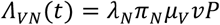

and

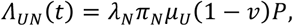

respectively, assuming vaccination does not impact susceptibility to the test-negative conditions.

Test-negative studies typically measure the OR of vaccination among the test-positive and test-negative subjects, similar to the exposure OR in case-control studies, using cumulative cases (*C*). For the test-positive outcome,

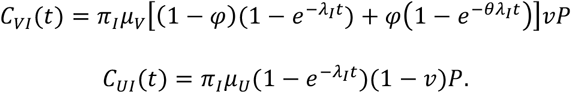

Under the assumption that test-negative infections are not immunizing, cumulative cases are proportional to the incidence rate and study duration:

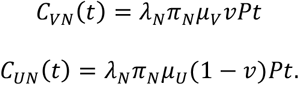

We consider the case of immunizing test-negative outcomes in **Web Appendix 1**. Using the vaccine-exposure OR measured from cumulative cases,

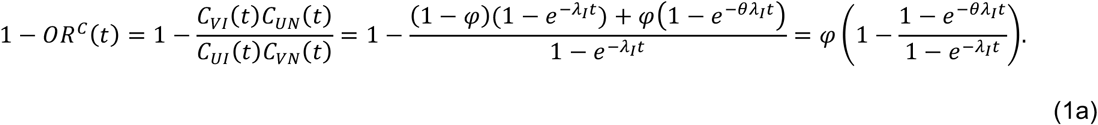

Under the special case of “all-or-nothing protection” (*θ*=0),

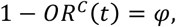

equal to the vaccine direct effect against infection. In contrast, under the special case of “leaky” protection for all recipients (*φ*=1),

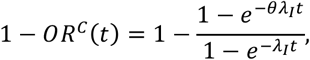

resulting in a bias toward the null value of 0. This bias is nonexistent near 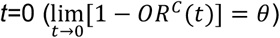, but grows as *t* increases 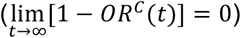.

Despite the lack of data in test-negative studies on the population (or person-time) at risk for infection, this result (eq. 1a) is equal to VE measures from the relative risk in randomized controlled trials. While methods have previously been proposed to recover the vaccine effect on susceptibility through uses of population-at-risk or person-time-at-risk data (23,29), we note that the absence of such measurements presents a unique obstacle to bias correction in test-negative studies.

Studies may also measure time-specific ORs, for instance by stratifying analyses into sub-seasonal intervals (30–33) or by interacting vaccination and time in logistic regression models fitted to individual-level data (34,35). In comparison to ORs estimated from cumulative cases, such estimates are often sought to gauge differences over time in VE, for instance due to waning of protection. As the time increment approaches zero, terms included in the OR approach the ascertainment rates of test-positive and test-negative subjects. We therefore define this measurement as

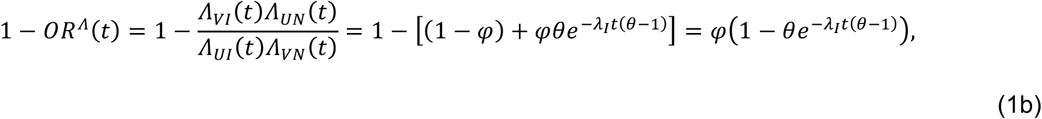

again reducing to

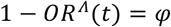

under “all-or-nothing” protection but allowing bias to persist under “leaky” protection for all recipients

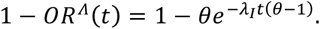

Here bias is again nonexistent at *t*=0 and worsens as *t* → ∞, further increasing with *λ_I_*. Intuitively, the bias arises due to differential depletion of vaccinated and unvaccinated susceptible individuals, consistent with other study designs (23,36,37). Presuming the vaccine is efficacious, more unvaccinated than vaccinated individuals will have been depleted later in the epidemic, confounding instantaneous comparisons.

We illustrate functional forms of 1 − *OR^c^*(*t*) and 1 − *OR^Λ^*(*t*) under scenarios of “leaky” and “leaky-or- nothing” protection in **Figure 1** and **Figure 2**, respectively. Considering first a “leaky” vaccine with 1 − *θ* = 50% efficacy, under conditions of *λ_I_*=0.001, 0.005, and 0.01 infections/person-day, VE estimates based on cumulative cases are 2.2%, 11.2%, and 22.1% lower than the true vaccine efficacy, respectively, after 90 days of influenza transmission (**Figure 1**); by this point, 8.6%, 36.7%, and 59.3% of unvaccinated individuals are expected to have been infected. Serological studies have revealed cumulative infection rates in the range of 20-40% for seasonal influenza (38,39) and up to 47% for influenza A(H1N1)pdm09 (40) among unvaccinated (and presumably susceptible) children, suggesting reported VE estimates may fall in the middle of this range in terms of bias; differences in susceptibility across ages and risk strata may, however, result in differential rates of infection and differential degrees of bias in estimates (41). The exposure-dependent biases we identify worsen with lower vaccine efficacy: for 1 − *θ* = 20%, estimated values fall 3.6%, 17.1%, and 32.4% below the true effect for *λ_I_*=0.001, 0.005, and 0.01, respectively. Estimates based on the ascertainment rate (1 − *OR^Λ^*(*t*)) would show greater bias at the same point in time: VE estimates are reduced by 4.6%, 25.2%, and 56.8% for a vaccine conferring 50% efficacy, and by 7.3%, 37.7%, and 78.9% for a vaccine conferring 20% efficacy.

**Figure 1:**
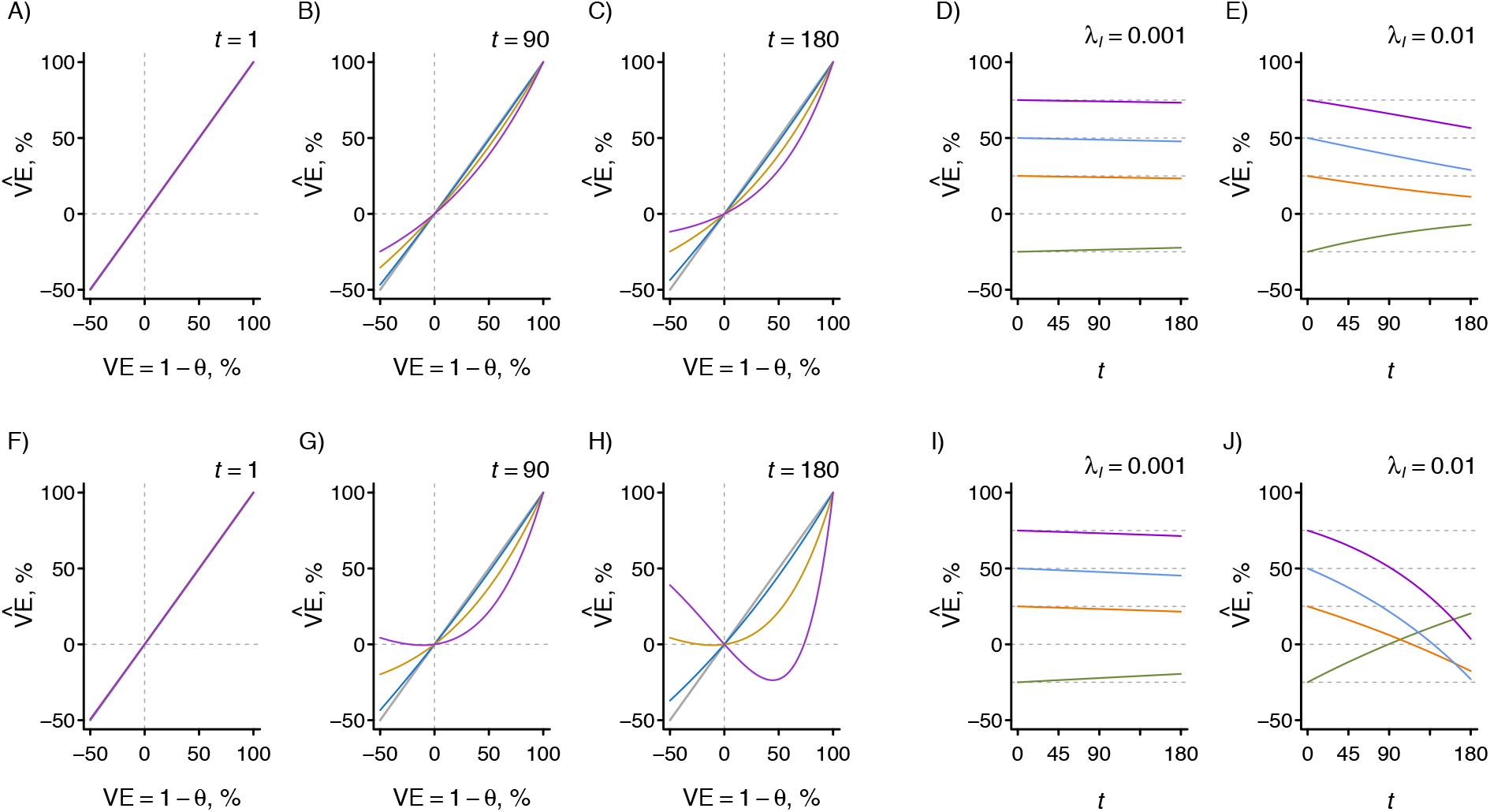
Test-negative measures under “leaky” protection. We illustrate test-negative VE estimates obtained from the exposure odds ratio for a vaccine conferring “leaky” protection (*φ*=1) to all recipients; Figure 2 includes extensions to “leaky-or-nothing” protection with differing values of *φ*. Estimates use (A–E) cumulative case data (1 − *OR^c^*) and (F–J) ascertainment rates (1 – *OR^Λ^*), under an assumption of no correlation between vaccination and exposure or susceptibility. Panels A–C and F–H illustrate measurements at set times (*t*) under differing transmission intensity (*λ_I_* equal to rates of 0.001, 0.005, and 0.01 infections per susceptible day at risk for blue, orange, and purple lines, respectively). Panels D, E, I, and J illustrate changes over time in estimated vaccine effectiveness, under scenarios of vaccine effectiveness equal to −25%, 25%, 50%, and 75% for green, orange, blue, and purple lines, respectively.

**Figure 2:**
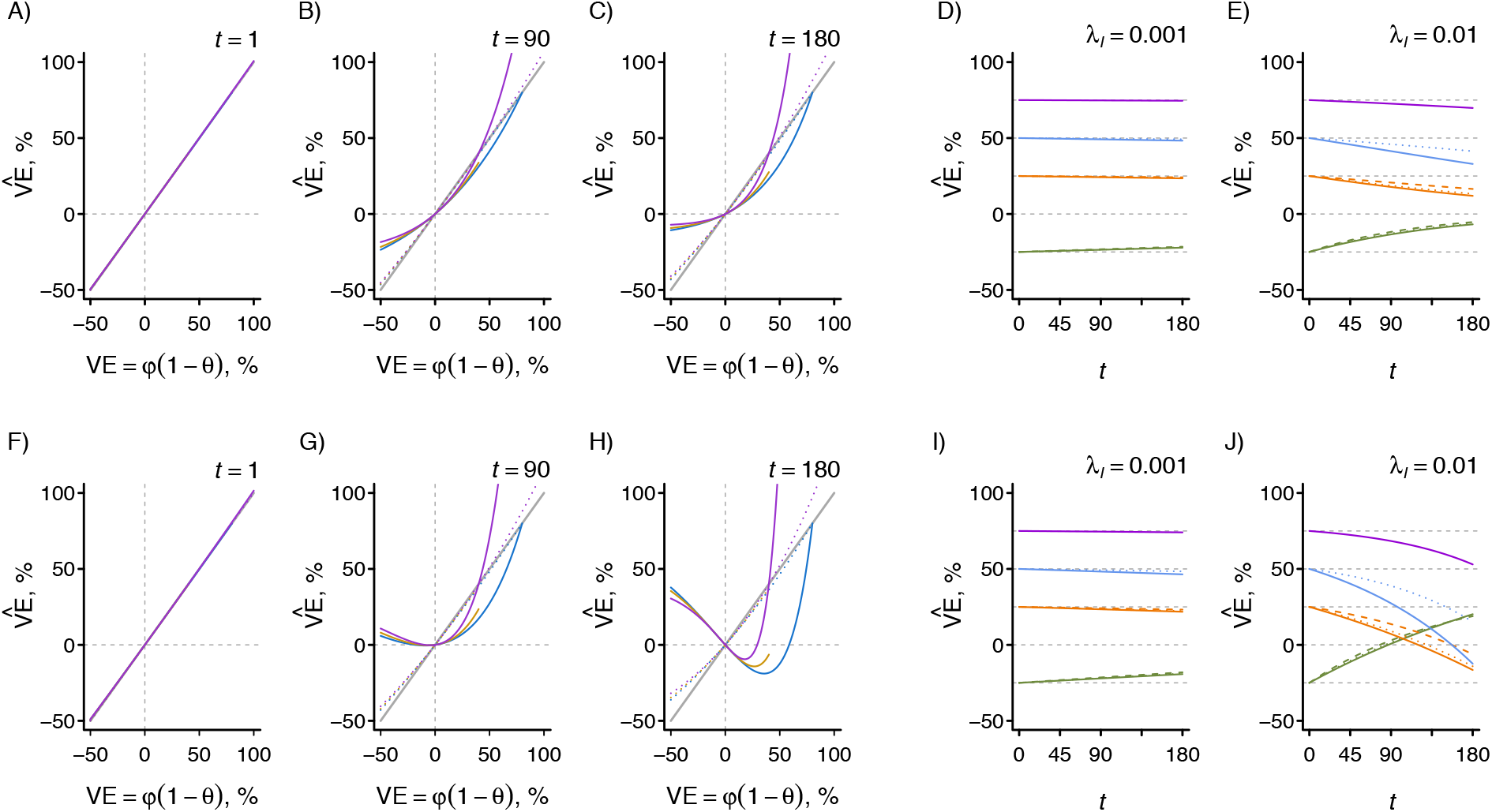
Test-negative measures under “leaky-or-nothing” protection. We illustrate test-negative VE estimates obtained from the exposure odds ratio for a vaccine conferring “leaky-or-nothing” protection; compare against Figure 1 for the special case of “leaky” protection (*φ*=1). Estimates use (A–E) cumulative case data (1 − *OR*^C^) and (F–J) ascertainment rates (1 − *OR^Λ^*), under an assumption of no correlation between vaccination and exposure or susceptibility. Panels A–C and F–H illustrate measurements at set times (*t*) under differing transmission intensity (*λ_I_* equal to rates of 0.001 and 0.01 infections per susceptible day at risk for dotted and solid lines, respectively); we illustrate performance of the estimator with differing degrees of vaccine response, illustrating *φ* equal to 0.8, 0.6, and 0.4 for blue, orange, and purple lines, and *θ* = (*φ − VE*)*φ*^−1^. Panels D, E, I, and J illustrate changes over time in estimated vaccine effectiveness. As in Figure 1, green, orange, blue, and purple lines signify scenarios of −25%, 25%, 50%, and 75% vaccine effectiveness; dashed, dotted, and solid lines signify *φ* equal to 0.4, 0.6, and 0.8, respectively.

Figure 2 illustrates how bias is further influenced by the contributions of vaccine response probabilities to overall vaccine efficacy. For a vaccine conferring 50% efficacy based on 90% of individuals responding (so that 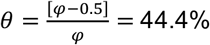), and again defining *λ_I_*=0.001, 0.005, and 0.01 infections/person-day, 1 − *OR^c^*(*t*) as of *t*=90 yields values subject to 2.0%, 10.0%, and 20.0% downward bias, respectively. With the same efficacy based on 60% of individuals responding (*θ*=16.7%), the degree of bias is reduced to 0.8%, 3.9%, and 8.2% below the true effect.

To aid interpretation in the context of previous studies (2,11), we also illustrate the modeled causal process using a directed acyclic graph (**Figure 3**), revealing that the special case of “all-or-nothing” protection precludes bias from vaccine-derived protection against influenza infections occurring before the ARI episode for which an individual seeks care. We derive VE estimators accounting for additional real-world circumstances—including time-varying transmission intensity during an influenza season and the use of naturally immunizing test-negative endpoints—in **Web Appendix 1**, showing that the OR retains the biases identified under our simpler initial assumptions.

**Figure 3:**
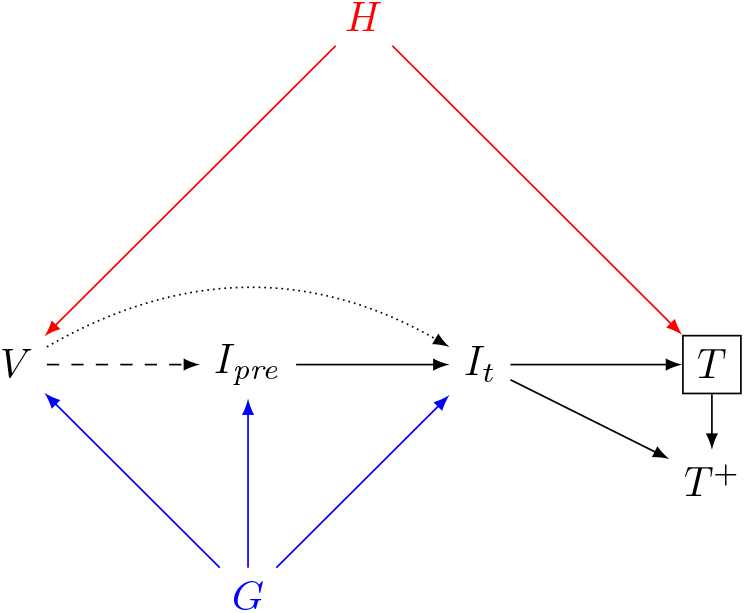
Causal directed acyclic graph (DAG) illustrating a key source of bias for leaky vaccines. Healthcare seeking (H) drives receipt of the vaccine (V) as well as receipt of a test (T). By design, studies select on testing, as only tested individuals are included. The effect of interest, signified by the dotted arrow, is that of vaccination on influenza at the time of testing (I*_t_*). However, influenza may also occur at a preceding point in the season (I_pre_, dashed arrow). The test-positive outcome (T^+^) arises when an individual is infected at the time of testing (I*_t_*→T^+^). Natural immunity prevents influenza re-infection during the season (I_pre_→I*_t_*). The fact that I_pre_ is not—and cannot be—conditioned on leads to a second pathway not of direct interest (V→I_pre_→I*_t_*) biasing the estimate of the direct effect V→ I*_t_* in the leaky-vaccine case. This bias is not present in the case of all-or-nothing protection. Here, two distinct subgraphs can be envisioned. In the first, applicable only to the proportion (*φ*) of protected, vaccinated individuals, the path V→>I_pre_→>If is not of concern, as Pr[I_pre_|V]=0. In the second, applying to the remaining proportion (1−*φ*) of unprotected individuals, the paths V→I_pre_→I*_t_* and V→I*_t_* are null, consistent with the situation where V=0. Subsequent sections of this manuscript focus on other sources of bias evident in this DAG. The first concerns the impacts of a confounder (G) of exposure or susceptibility to influenza infection (here indicated in blue; see also Figure 3 of Lipsitch and colleagues, 2017 (2)); the second concerns selection bias resulting from differential healthcare-seeking behavior among vaccinated and unvaccinated persons along the pathway V←H→T (here highlighted in red).

### Conditions for sign bias

In some applications, testing for a protective or harmful effect of the vaccine may take priority over obtaining precise measurements of effect size. The conclusions of such hypothesis tests rest on an assumption that the OR is not subject to sign bias, reflecting the circumstance *OR*(*t*) > 1 for an effective vaccine (as defined by the condition *θ* < 1), or *OR*(*t*) < 1 for an ineffective vaccine (for which *θ* > 1). The plots of 1 − *OR^Λ^*(*t*) in **Figures 1 and 2** illustrate that VE estimates based on the ascertainment rate of cases may encounter sign bias. The OR measured from ascertainment rates reaches one—suggesting no vaccine effect—when the cumulative transmission to which a population has been exposed (*λ_I_×t*) reaches a particular threshold:

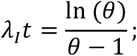

we derive this threshold in **Web Appendix 2**. Once at least this proportion of the unvaccinated population has become immune to infection, cases will appear with higher frequency among vaccinated than unvaccinated individuals even when vaccine-derived protection does not wane. These circumstances demonstrate the need for caution in interpreting time-specific (continuous or sub-seasonal) VE measurements from test-negative studies (30–33), or for strategies to account for previous infection prevalence among vaccinated and unvaccinated persons.

## PERFORMANCE OF THE ODDS RATIO UNDER DIFFERENTIAL EXPOSURE OR SUSCEPTIBILITY OF VACCINATED AND UNVACCINATED PERSONS TO THE CONDITIONS

The test-negative design is typically employed in observational studies where individuals have received vaccination voluntarily. In contrast to assumptions in the above section that vaccination is uncorrelated with exposure or susceptibility to infection, variation in vaccine uptake across risk groups is well-recognized (2). For instance, preferential vaccine receipt has been reported among relatively healthy older adults (42,43) and among persons prioritized for vaccination such as healthcare workers (who may have elevated risk of encountering infected persons) and individuals with underlying health conditions (who may be at risk for severe outcomes if infected) (44,45). This circumstance corresponds to the presence of a confounder (“G” in **Figure 3**) related to disease risk as well as vaccination.

Absent vaccine-derived protection, define *α_VI_*/*α_UI_* and *α_VN_*/*α_UN_*> as the relative rates at which individuals who seek vaccination would be expected to acquire influenza and test-negative conditions, respectively, measured against the expected rates among individuals who do not seek vaccination. These relative rates do not consider the biological effect of the vaccine, but only the counterfactual associated with vaccine-seeking status.

Accounting further for vaccine-induced protection, the ascertainment rates of test-positive and test-negative subjects are

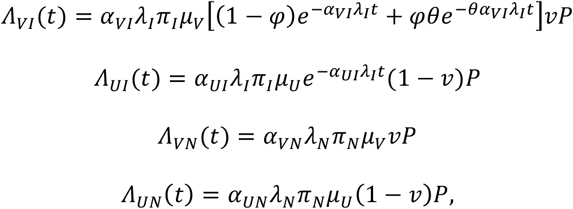

resulting in cumulative case measures

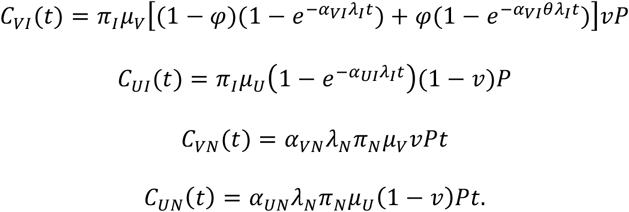

Estimating VE from cumulative cases,

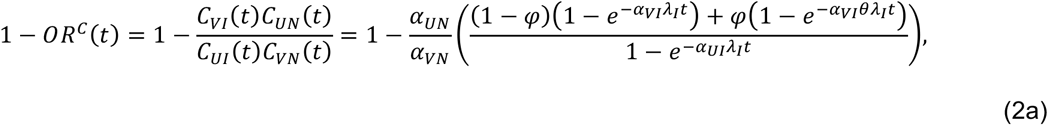

whereas the estimate based on ascertainment rates is

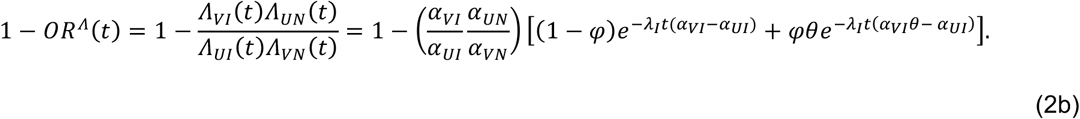

These estimates reduce to

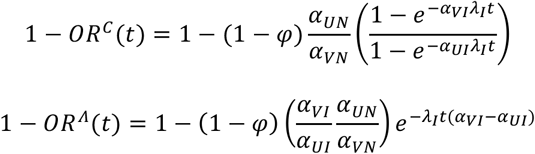

under “all-or-nothing” protection, and

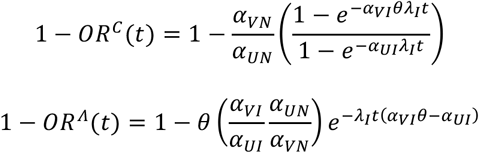

under “leaky” protection.

Consider alternatively that *π_VI_*/*π_UI_* and *π_VN_*/*π_UN_* are the relative risks of ARI given influenza and test-negative infections, respectively, for individuals who seek vaccination, measured against the risk among individuals who do not seek vaccination; we again distinguish that these differences owe to factors other than vaccine-derived protection (17), and consider vaccine protection against disease progression in a subsequent section. Incorporating *π_I_* and *π_N_* into the ORs formulated above,

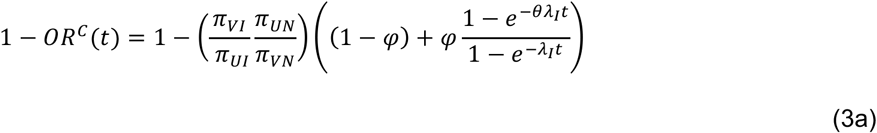

and

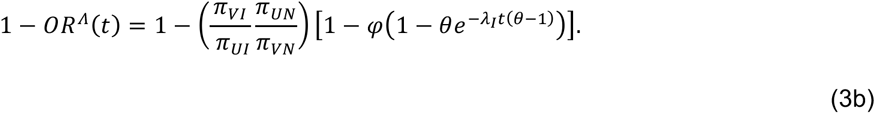

Under “all-or-nothing” protection,

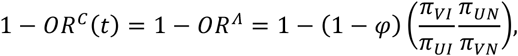

reducing to *φ* if differences between vaccinated and unvaccinated persons equally affect progression of influenza and test-negative conditions to symptoms, i.e. 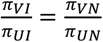. For a vaccine conferring “leaky” protection to all recipients,

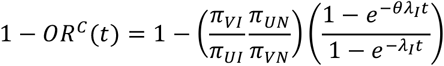

and

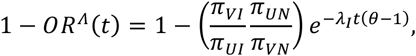

reducing when 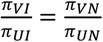 to the bias present when vaccine-seeking is uncorrelated with exposure or susceptibility to infection (eqs. 1a and 1b).

Incorporating heterogeneity in both acquisition and progression,

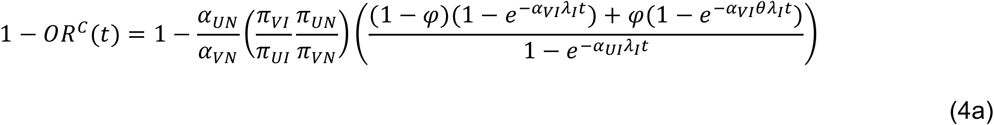

and

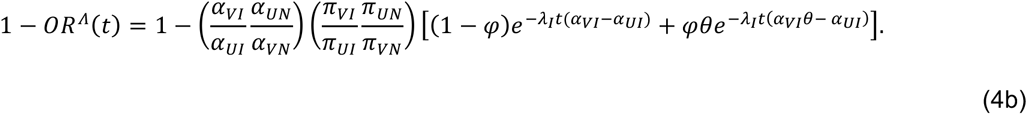

These circumstances underscore that differential vaccine uptake among persons at high and low risk for infection or for symptoms given infection—a well-known phenomenon in observational studies of vaccines and other health interventions—may undermine causal interpretations of the OR in test-negative studies.

## BIAS ASSOCIATED WITH DIFFERENTIAL TREATMENT SEEKING AMONG THE VACCINATED AND UNVACCINATED

To this point we have considered ARI as a singular clinical entity and assumed all individuals seeking care for ARI are tested for influenza. However, different infections may cause clinically-distinct presentations influencing the likelihood that individuals seek treatment, or the likelihood that clinicians test for influenza (46). Here we address the possibility for such a scenario to lead to selection bias from conditioning on the collider T (testing), the pathway 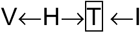 in **Figure 3**.

Consider that the spectrum of clinical presentations can be discretized into “moderate” (*M*) and “severe” (*S*) classes, occurring with probabilities 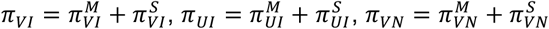, and 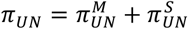. Define 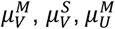, and 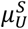 as the associated probabilities of seeking care given symptoms and vaccination status, and let *ξ^M^* and *ξ^S^* indicate the probabilities of receiving a test given symptoms. Bias associated with differential treatment-seeking persists unless the relative risk of testing given infection (which includes experiencing symptoms, seeking treatment, and being tested) does not differ for influenza and other conditions:

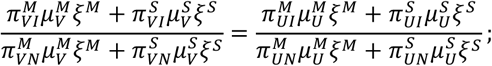

we derive the associated VE estimators in **Web Appendix 3**. Expressed more generally, this bias arises unless

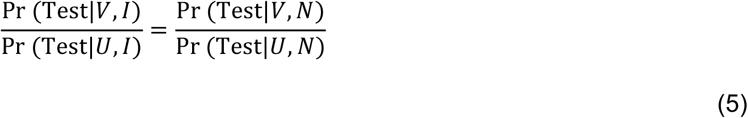

when accommodating all possible factors that influence whether individuals are tested. Ensuring that the above condition is met can guide study implementation and circumvent possible biases owing to associations of vaccination with care-seeking given illness, receipt of clinical testing, and willingness to participate in the study.

### Correction of bias through the use of clinical criteria for enrollment and testing

A possible correction exists when enrollment and testing are tied to stringently-defined clinical criteria, i.e. criteria for which eq. 5 holds. For example, if tests are performed conditioning on cases resembling a well-defined and monotypic “Severe” entity (substituting *ξ^M^*=0 in eqs. 5a and 5b), the OR retains bias only from differential infection rates and symptom risk between the vaccinated and unvaccinated:

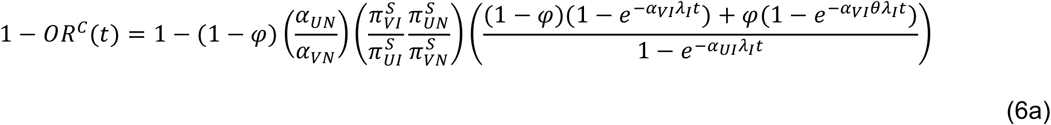

when measured from cumulative incidence, or

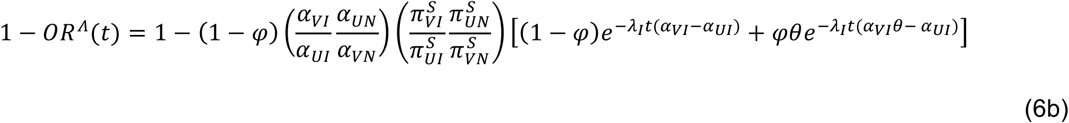

when measured from the ascertainment rate (resembling eqs. 4a and 4b). Absent any association of the decision to receive the vaccine with individuals’ exposure or susceptibility to infection and ARI, eqs. 6a and 6b reduce to eqs. 1a and 1b.

## MEASURING VACCINE EFFECTIVENESS AGAINST PROGRESSION

In addition to protection against infection, reductions in symptom risk given infection are of interest in VE measures (17). Define *ρ* as the relative risk for vaccine-protected individuals to experience symptoms given infection owing to vaccine-derived immunity. When decisions to vaccinate are not correlated with exposure or susceptibility to the infections, other than through vaccine-derived immunity,

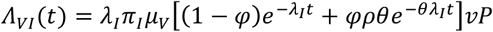

and

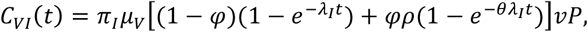

so that

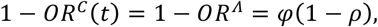

Under the special case that a vaccine reduces risk of symptoms without protecting against infection (*θ*=1)—as might apply to oral cholera vaccines (47–49)—these measures reduce to

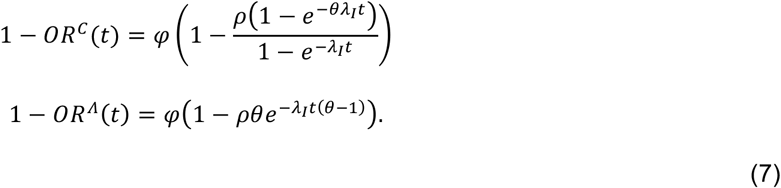

an unbiased estimate of VE against progression. Under confounding between vaccination and exposure or susceptibility to the infections,

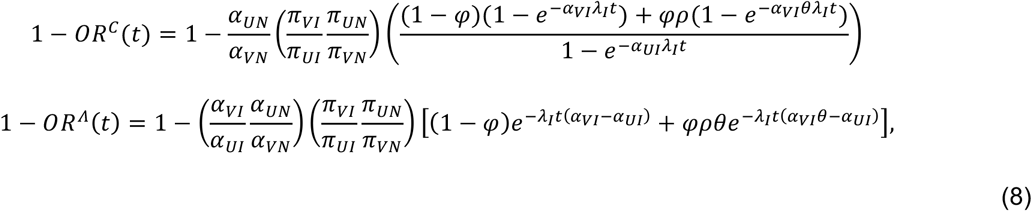

reducing to

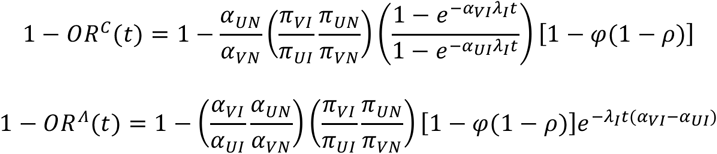

for a vaccine protecting against symptoms only (*θ*=1).

## IMPLICATIONS

Recent years have seen growing enthusiasm about the integration of data from observational studies in decisions surrounding influenza vaccine policy (50), in part based on belief that vaccine direct effects— which have traditionally been measured in prospective, randomized controlled trials—can be recovered under the test-negative design (6,10,16,24). However, uptake of the test-negative design by researchers and policymakers has preceded thorough examination of its theoretical justification (14). Our analysis highlights limitations to interpreting VE estimates based on the exposure OR from test-negative studies.

Our most troubling finding is that the OR measured by test-negative studies is unsuited to estimating the vaccine direct effect on susceptibility to infection even under circumstances consistent with randomized vaccine allocation, unless protection is known to follow an “all-or-nothing” mechanism of action. These results echo longstanding concerns about measurement of the effectiveness of “leaky” vaccines in case-control studies (23,51–53) as well as clinical trials (36,37), and occur because unvaccinated persons become immune via natural infection faster than unvaccinated ones. Researchers rarely know *a priori* to what extent a vaccine confers “leaky” or “all-or-nothing” protection, making it difficult to know under what circumstances studies may be subject to the resulting bias.

We also show that certain traditionally-recognized sources of confounding in observational studies— arising due to differential exposure or susceptibility to infection and symptoms among vaccinated and unvaccinated persons—persist under the test-negative design. Because resulting biases may lead to time-varying estimates of VE, declines in 1-OR over a season may not support inference of waning vaccine protection (31–35). Last, whereas the test-negative design has been viewed as a strategy to eliminate treatment-seeking bias, we find that bias may persist under differential symptom severity for influenza and test-negative infections.

Several assessments of test-negative studies based on DAGs (2,11) have pointed to similar sources of confounding, and the practical importance of these findings has been debated amid uncertainty about the magnitude of associated bias in estimates (16). The framework we have taken provides a basis for quantifying bias directly. We show that the OR of test-negative studies can supply VE estimates that are not equal to the causal vaccine effect on susceptibility and that sign bias may arise such that the instantaneous OR leads to incorrect inferences about whether a vaccine is effective or not. This is contrary to the frequent assumption that the OR provides, at minimum, a valid and direction-unbiased test of the null hypothesis of no causal effect (11).

Other approaches have been taken to assess bias in test-negative studies. In informal comparisons, VE estimates from test-negative studies of live oral rotavirus vaccines and oral cholera vaccines have appeared similar to vaccine efficacy estimates from randomized controlled trials in the same settings (54,55). While these findings may suggest the biases we identify are not always large in practice, our study and others (36,37) have pointed to potential sources of bias that may also affect estimates of the vaccine direct effect in randomized controlled trials. Moreover, seasonal influenza vaccine trials are not conducted on a year-to-year basis amid alterations to the strain composition of vaccines and changes to the immune profile of hosts. This has led to difficulty accounting for instances where conclusions of randomized controlled trials and test-negative studies have appeared to be in conflict. For instance, LAIV effectiveness has appeared poor in test-negative studies undertaken since the emergence in 2009 of a novel H1N1 influenza A virus (56,57), despite superior efficacy of LAIV over inactivated influenza vaccine among children in earlier randomized controlled trials (58–60).

Many of the biases we identify result from differential acquisition of natural immunity among vaccinated and unvaccinated persons. The strength and duration of such immunity differs among infectious diseases for which test-negative studies have been undertaken to estimate VE; specific implications for weakly-immunizing infections such as rotavirus (18,19) and respiratory bacterial agents (4,22) should be assessed. Uses of the test-negative design in increasingly innovative applications, for instance an evaluation of cluster-randomized deployments of Wolbachia-infected mosquitoes to prevent dengue (61,62), further merit consideration in terms of transmission dynamic parameters such as we consider here.

## STRATEGIES TO COUNTERACT BIAS

While our analysis identifies limitations to the validity of VE estimates based on the vaccine-exposure OR under the test-negative design, the results highlight specific improvements that can be made to the interpretation of data from test-negative studies. We have shown that the use of strict clinical criteria or case definitions for enrollment and testing can reduce bias due to differential healthcare-seeking behavior among vaccinated and unvaccinated persons. Whereas test-negative studies typically stratify estimates according to influenza type/subtype or even the genetic clade, our findings suggest bias may persist if there are meaningful epidemiologic differences in risk factors for infection and disease among vaccinated and unvaccinated persons. This bias can be reduced by stratifying estimates to minimize within-stratum differences in exposure or susceptibility to infection among vaccinated and unvaccinated persons. While we point out the inability of test negative studies to measure “leaky” or “leaky-or-nothing” protection accurately, the persistence of such bias in randomized controlled trials echoes a broader need to consider epidemiological approaches for the measurement of imperfect forms of immunity (23,51–53). Because biases resulting from the “leaky” nature of vaccine protection have lower impact in populations less exposed to transmission, VE estimates from early in the influenza season may be more reliable than those obtained later. This circumstance suggests a need to maximize statistical power for test-negative studies in the initial weeks or months of seasonal or pandemic influenza transmission. In addition, monitoring the cumulative incidence of infections in populations, for instance through serological studies, could facilitate correction for the differential prevalence of naturally-acquired immunity among vaccinated and unvaccinated persons. Evidence from test-negative studies of VE against influenza should be interpreted with the limitations we report here in mind, in particular for vaccination policymaking.

## ACKNOWLEDGMENTS

Author Affiliations: Division of Epidemiology, School of Public Health, University of California, Berkeley, Berkeley, California (Joseph A. Lewnard); Center for Communicable Disease Dynamics, Department of Epidemiology, Harvard TH Chan School of Public Health, Boston, Massachusetts (Joseph A. Lewnard, Christine Tedijanto, and Marc Lipsitch); and WHO Collaborating Center for Infectious Disease Epidemiology and Control, School of Public Health, Li Ka Shing Faculty of Medicine, The University of Hong Kong, Hong Kong Special Administrative Region, China (Benjamin J. Cowling).

This work was supported by National Institute of General Medical Sciences grant U54GM088558.

## FOOTNOTES PAGE

### Abbreviations

ARI: : Acute respiratory illness
DAG: : Directed acyclic graph
LAIV: : Live attenuated influenza vaccine
OR: : Odds ratio
VE: : Vaccine effectiveness (used interchangeably with vaccine efficacy in this context as the estimand of both observational and randomized studies, and defined as the causal effect of the vaccine on susceptibility of individuals to infection and/or disease)

## WEB APPENDIX 1

Here we derive estimators maintaining the assumption of vaccination unconfounded by exposure or susceptibility to the conditions to consider the impact of time-varying (seasonal) infection rates and the use of immunizing test-negative conditions.

### Time-varying infection rates

Consider that transmission intensity varies over time, so that acquisition rates are *λ_I_*(*τ_j_*) at time *τ_j_*, consistent with the seasonal rise and fall of influenza transmission. Indexing by day, the probability of evading infection to time *t* is 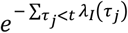, which can be substituted for *e^−λ_I_t^* in eqs. 1a and 1b so that

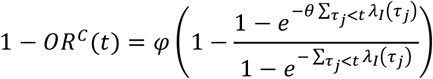

and

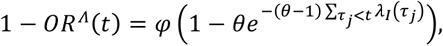

resembling the expression for time-invariant *λ_I_* and retaining the relevant biases, which scale with cumulative exposures transmission over time 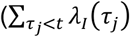 in place of *λ_I_×t*).

### Immunizing test-negative conditions

Under an assumption that test-negative conditions are selected which lend protective immunity after infection, preventing reacquisition,

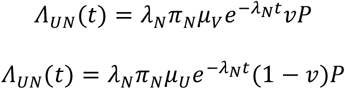

and

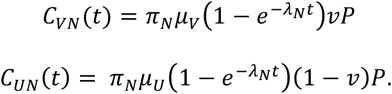

The terms describing the cumulative proportions of vaccinated and unvaccinated persons infected by the test-negative condition (exp[−*λ_N_t*]), and the proportions of vaccinated and unvaccinated persons remaining susceptible (1−exp[−*λ_N_t*]), cancel in the expressions for *OR^c^*(*t*) and *OR^Λ^*(*t*), respectively, which invoke (1−exp[−*λ_N_t*])/(1−exp[−*λ_N_t*]) and exp[−*λ_N_t*]/exp[−*λ_N_t*], respectively. Thus, our original VE derivations apply to the scenario of immunizing test-negative conditions.

## WEB APPENDIX 2

Here we derive the conditions under which the OR may inaccurately convey the protective (or risk-increasing) effect of vaccination, assuming no correlation of the decision to vaccinate with exposure or susceptibility to infection and disease. Consider the condition

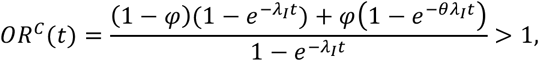

which implies

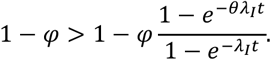

and thus *θ* > 1. Sign bias is present if *θ* < 1, so that the necessary conditions for sign bias cannot be met. In the converse situation, *OR^c^*(*t*) < 1 implies *θ* < 1, which again cannot be true when *θ* > 1. Thus, sign bias does not occur in measurements based on cumulative cases, provided vaccination is uncorrelated with exposure or susceptibility to the infections. Because such confounding is likely in real-world studies, we assess resulting biases in a later section.

Figure 1 and Figure 2 illustrate that sign bias may affect measurements based on the ascertainment rates of test-positive and test-negative subjects. For *θ* < 1, the condition

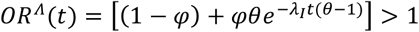

implies

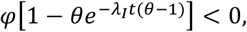

so that sign bias arises when

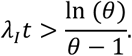

Conversely, for *θ* > 1, sign bias (indicated by *OR^Λ^*(*t*) < 1) arises under

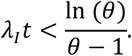

## WEB APPENDIX 3

Here we derive the OR under a scenario of differential severity of ARI caused by test-positive and test-negative conditions. Considering “moderate” and “severe” classes of symptomatic infection, ascertainment rates of subjects are

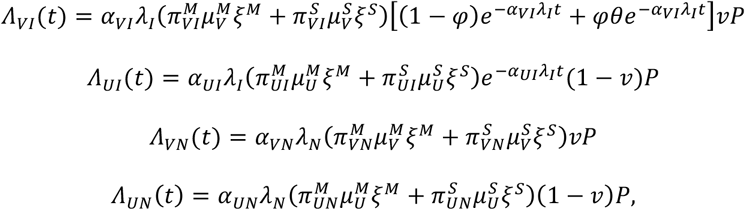

so that

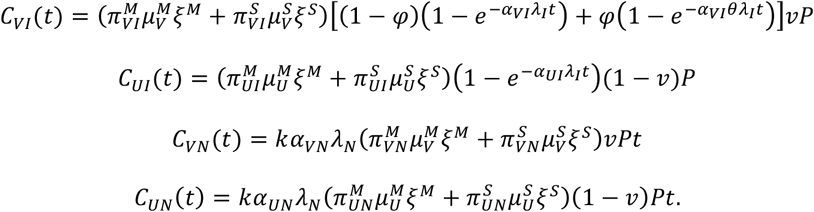

The test-negative VE measures reduce to

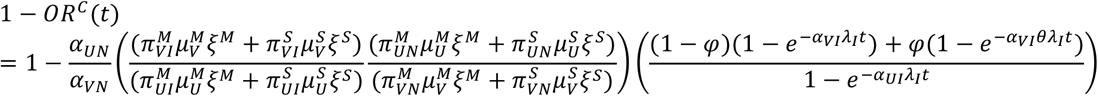

and

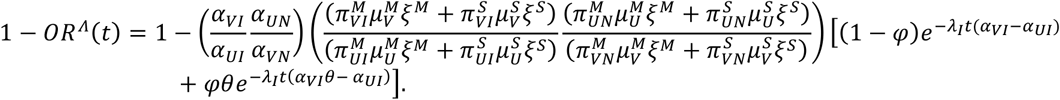

